# Identification of Alkaloids and Related Intermediates of *Dendrobium officinale* by Solid-Phase Extraction Coupled with High-Performance Liquid Chromatography Tandem Mass Spectrometry

**DOI:** 10.1101/2022.02.09.479720

**Authors:** Cheng Song, Yunpeng Zhang, Muhammad Aamir Manzoor, Guohui Li

## Abstract

JA signaling plays a pivotal role in plant stress responses and secondary metabolism. Many studies have demonstrated that jasmonates effectively induce the expressions of alkaloid biosynthesis genes in various plants, which rendered to the accumulation of alkaloid to counteract stresses. Despite the multiple roles of jasmonate (JA) in the regulation of plant growth and different stresses, less studied involved in the regulatory role of JA in *D. officinale* alkaloids. A strategy for the rapid identification of alkaloid and the intermediates of *D. officinale* was established based on a solid-phase extraction coupled with high-performance liquid chromatography tandem mass spectrometry method. By using SPE-HPLC-LTQ-Orbitrap method, the potential compounds were tentatively identified by aligning the accurate molecular weight with the METLIN and Dictionary of Natural Products databases. The chemical structures and main characteristic fragments of the potential compounds were further confirmed by retrieving the multistage mass spectra from the MassBank and METLIN databases.The Mass Frontier software was used to speculate the fragmentation pathway of the identified compounds. Seven alkaloids were separated and identified from *D. officinale*, which were mainly classified into five types (tropane alkaloids, tetrahydroisoquinoline alkaloids, quinolizidine alkaloids, piperidine alkaloids and spermidine alkaloids). Besides the alkaloids, forty-nine chemical substances, including guanidines, nucleotides, dipeptides, sphingolipids and nitrogen-containing glucosides, were concurrently identified. These findings gives the composition of chemicals currently found in *D. officinale*, which could provide the scientific method for the identification of alkaloids in other *Dendrobium* plants.

## Introduction

Traditional Chinese medicine (TCM) involves complex chemical constituents. An effective determination of one group of metabolites from a certain plant entails a comprehensive methodology for their purification and identification. Conventional detection methods such as thin-layer chromatography (TLC) and high-performance liquid chromatography (HPLC) generally provide such low accuracy and poor sensitivity that leading to incapable of detecting some trace substances (Pan et al., 2015). A separation and identification method with strong specificity, high efficiency, and few interference is urgently needed. Solid-phase extraction (SPE) was usually used for the complicated specimens by reducing matrix interference and raising sensitivity (Bucar et al., 2013). Orbitrap mass spectrometry is one kind of high-resolution mass analyzer containing high-resolution multistage mass fragment monitoring, a selective fragmentation mode, and a personalized data acquisition method (Qiu et al., 2015). Orbitrap has been widely applied in pharmacokinetic analysis, food industry, environmental monitoring and proteomics (Quifer-Rada et al., 2015; Li et al., 2013). Therefore, the SPE-HPLC-LTQ-Orbitrap technique is proved to be a powerful means to elucidate the complicacy in TCM. *D. officinale*, as a type of perennial herb, is attached to the Dendrobium genus of Orchidaceae. As a valuable TCM, *D. officinale* contains many different types of active ingredients, including polysaccharides, alkaloids, bibenzyls, phenanthrenes, sesquiterpenoids, fluorenones, flavonoids, steroids, *etc*. (Xu et al., 2013). Some of these compounds had exhibited versatile pharmacological and pharmacodynamic activity (Ng et al., 2012). The earliest study on the alkaloids of the Dendrobium genus started from dendrobine of *D. nobile*. For the decades, dozens of alkaloids have been subsequently identified from *D. anosmum, D. chrysanthum, D. crepidatum, D. findlayanum, D. friedricksianum, D. hilderbrandii, D. loddigessi, D. lohohense, D. moniliforme, D. parishii, D. pierardii, D. primulimun, D. snowflake* and *D. wardianum* (Xu et al., 2013). At present, the alkaloids identified from Dendrobiun plants had been divided into five categories based on their chemical structures, including sesquiterpenoid alkaloids, indolizidine alkaloids, pyrrolidine alkaloids, phthalide alkaloids, and imidazole alkaloids (Song et al., 2022).

Thus far, the composition and biosynthetic pathway of the alkaloids in *D. officinale* has not been entirely understood. The determination of the alkaloids between *D. officinale* and *D. nobile* using an LC-MS method showed that 25 identical compositions in the two Dendrobium plants were tentatively identified; however, dendrobine was only detected in *D. nobile* (Song et al., 2020). Using a UPLC method, a few coumarins, alkaloids and bibenzyl components from fifteen Dendrobium species were quantitatively determined. Dendrobine was only detected in *D. nobile* originated from the Guizhou and Yunnan Provinces (Xu et al., 2010). The genome and transcriptome analyses of *D. officinale* implied that a biosynthetic pathway oriented to the terpenoid indole alkaloids likely exists (Guo et al., 2013; Yan et al., 2015). Recently, high-throughput sequencing revealed the identification of a total of 56 genes, including 25 key enzymes potentially involved in the synthesis of terpenoid indole alkaloids, tropine alkaloids and isoquinolines alkaloids in *D. officinale* (Niu et al., 2021). Many studies have shown that *CrMYC2* and *ORCAs* transcription factors play a positive role in elevating the TIAs content in *Catharanthus roseus* (Memelink et al., 2007). With the help of transcriptome sequencing, 75 bHLH family genes and 66 AP2/ERF genes were identified in the DEGs of *D. officinale*, suggesting that those transcription factors could be involved in the regulation of secondary metabolism, especially in TIAs biosynthesis (Zhang et al., 2016; Song et al., 2022). Mevalonate (MVA) and methylerythritol 4-phosphate (MEP) pathway are both core pathways related to MeJA-induced accumulation of TIAs (Song et al., 2021). In *D. officinale*, the key genes in MVA and MEP pathway and some post-modified enzymes were significantly up-regulated by MeJA treatment (Chen et al., 2019). In the previous study, high-yielding production of alkaloids from *D. officinale* were obtained with the combination of tryptophan, secologanin and MeJA treatment (Jiao et al., 2018; Song et al., 2022). Here, a technical scheme for the separation and determination of alkaloids and precursors in *D. officinale* was established. The accurate mass weights of the components separated by HPLC were determined using an Orbitrap mass analyzer. The multistage fragmentation mass spectra of alkaloids were monitored to speculate the potential substances based on a database-dependent retrieval method. Alkaloid and their precursors related to alkaloid-synthetic pathway were associated to construct the metabolic network between upstream and downstream. The results would provide a reliable scientific reference and technical guidance for the development and quality evaluation of *D. officinale*.

## Material and Methods

### Instruments and chemicals

The LTQ-Orbitrap XL mass spectrometer were manufactured by Thermo Fisher Scientific (Bremen, Germany). The Agilent 1260 instrument used in the experiments was manufactured by Agilent Technologies (Waldbronn, Germany). The Xcalibur (*v*. 2.1) and Mass Frontier (*v*. 6.0) were used for the analysis of mass spectrometry and speculation of the fragmentation pathways. Acetonitrile and formic acid were of HPLC grade. Other chemicals were domestically made analytical reagents. The standards of reserpine and sarpagine was obtained from Aladdin (Shanghai, China) and BioBioPha (Kunming, China), respectively.

### Plant samples preparation

The *D. officinale* samples used in the experiment were obtained from the tissue culture room of the Plant Cell Engineering Center in West Anhui University. The tissue-cultural seedlings cultivated in MS medium were collected after 100 μM MeJA treatment. The seedlings were firstly dried at 110 °C for 15 min and kept in a constant weight at 60 °C. The dried materials were ground into powders by a universal grinder. Using the ultrasonic extraction method, two gram of powder was placed in a round-bottom flask with 100 mL of methanol and extracted three times at room temperature. The bromocresol green colorimetric method was used for the determination of total alkaloids. The extraction solution was centrifuged for 10 min at 6000 x *g* to get the supernatant, then evaporated to dryness in a rotary evaporator. The extraction was dissolved with 5 mL of dilute sulfuric acid, filtered through a 0.22 μm membrane, and pretreated using MCX cartridges (60 mg, 3 mL) for purification. The protocol was successively conducted by our previous method (Song et al., 2020).

### LC-MS conditions

The Waters T3 column (150 mm × 4.6 mm, 3 μm) was used for the chromatographic separation. The mobile phases acetonitrile (A) with 0.1% formic acid and aqueous phase (B) with 0.1% formic acid were used for the experiment. Gradient elution process was as follows: 0-2 min (0% A), 2-12 min (0 to 15% A), 12-22 min (15 to 35% A), 22-32 min (35 to 80% A), and 32-37 min (80 to 0% A). The UV wavelength was set at 280 nm, and the column temperature was set at 25 °C. A three-way splitter was connected with the mass analyzer at a 0.3 mL min^-1^ flow rate. The ion-source parameters were refereed to Kumar’s method and slightly modified (Kumar et al., 2016). The detailed parameters were as follows: sheath gas at 25 arb, auxiliary gas at 3 arb, spray voltage at 4 kV, the capillary temperature at 320 °C, tube lens at 120 V, and capillary voltage at 30 V. The MS data were collected at 100 ≤ *m/z* ≤ 1000 in positive-ion mode. Two data acquisition methods were used in the experiments. In one method, a high-resolution scan was conducted using the Orbitrap mass analyzer to acquire the MS data at a resolution at 30,000 FWHM and the MS^2^ data at 15,000 FWHM. A data-dependent MS^n^ scan was used for the analysis of the MS^2^ spectra generated from the most abundant ions of the MS spectra. In the other method, a high-resolution scan was conducted first to acquire the MS data at a resolution at 30,000 FWHM, then the LTQ dynode was used to scan the MS^2^ and MS^3^ spectra. The dynamic exclusion function was used to reduce repeat scans, and the repeat count was 2. The exclusion duration was 20 s, and the exclusion mass width was 3 *m/z*. The suitable collision-induced dissociation (CID) energy was set at 35%. The minimum signal threshold was 500, and the isolation width of precursor ions was 2 *m/z*. Sarpagine and reserpine were respectively injected into the mass analyzer via a syringe pump at a flow rate of 6 μL min^-1^ to complete the direct infusion analysis. The putative molecular formulae were calculated and obtained according to the accurate molecular weight using Xcalibur (*v*. 2.1). Compounds were tentatively identified by matching METLIN, Mass Bank, Dictionary of Natural Products and Chemspider databases. For the compounds without multiple mass spectra, Mass Frontier (*v*. 6.0) was helped to speculate and deduce the fragmentation pathway.

## Results and discussion

### Biomass and alkaloid contents under the MeJA treatment

The effects of methyl jasmonate on the growth and alkaloid accumulation of *D. officinale* was studied. The results showed that MeJA delayed the growth rate of *D. officinale* and initiated the synthesis of a large number of alkaloids in logarithmic growth phase (**Figure 1**). The highest content of alkaloids climaxed to 343 μg/g when cultured on the 32 days. After that, the fresh weight tended to be stable, and the alkaloid content began to fall gradually. The results suggested that jasmonates could continuously promote the synthesis of alkaloids during the growth of *D. officinale*. The growth trend and accumulating pattern were consistent with our previous study (Jiao et al., 2018; Wang et al., 2021).

**Figure. 1.**
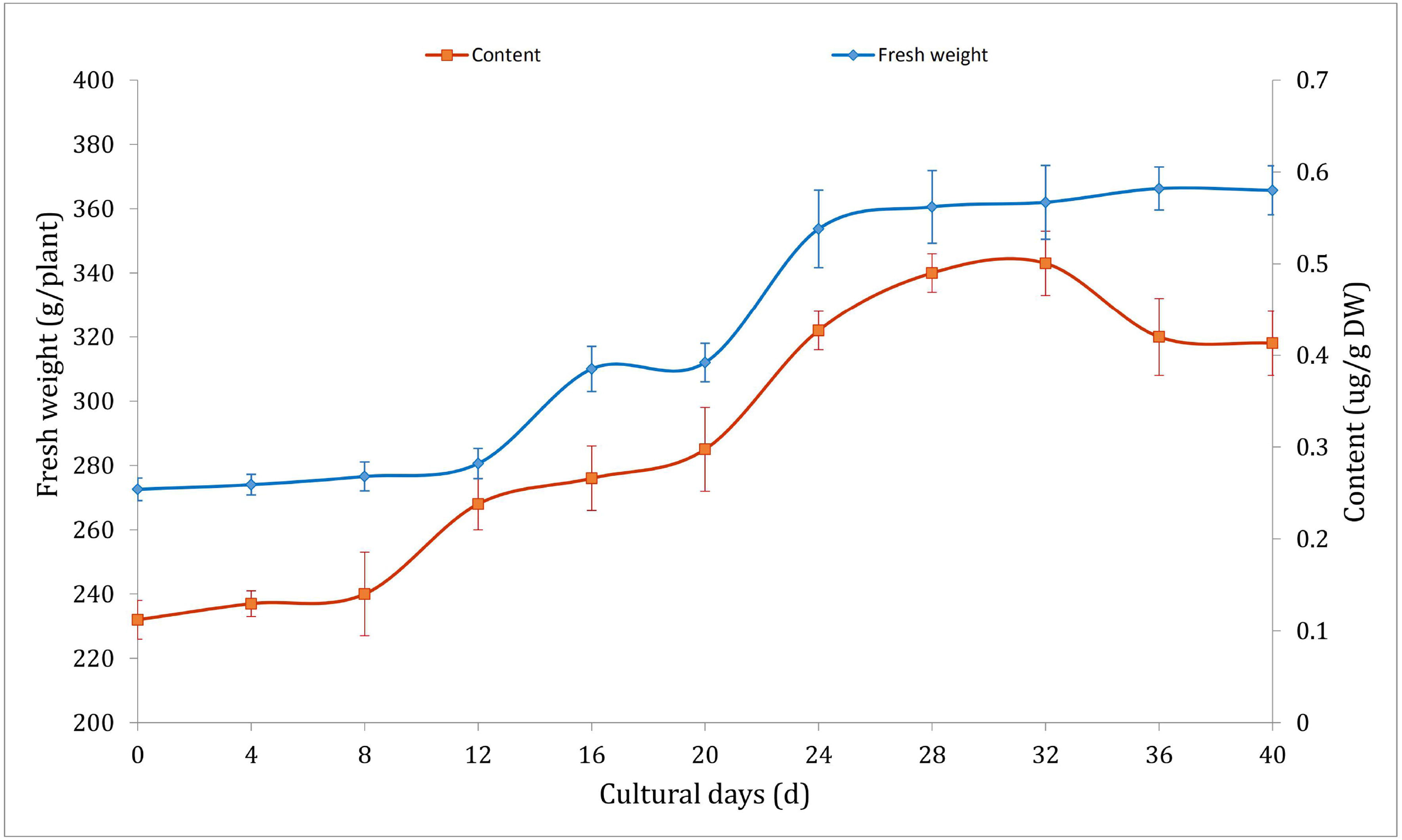
The fresh weight and alkaloid contents of *D. officinale* treated with 100 μM MeJA.

### Mass spectra interpretation

Firstly, *D. officinale* samples were extracted by methanol and dissolved with diluted sulfuric acid. The MCX cartridges were used for the purification of alkaloids from the acidic aqueous extraction. To optimize the condition of mass spectrometry, the suitable CID energy was achieved by the direct infusion experiment. Generally, the strongest fragment ions and the smallest number of fragments were observed at 30-35% collision energy, whereas the abundances of the fragments decreased substantially at 45% or greater collision energy. In this experiment, sarpagine and reserpine achieved sufficient fragments both in quantity and intensity at 40% collision energy. High-resolution mass analysis was conducted by a data-dependent MS^n^ scan from the workstation to produce abundant fragmentation spectra of the product ions. Due to the high resolution, the scan rate of a single spectrum was reduced, and the analytical analysis time of the entire sample was extended. An appropriate exclusion duration time was required for the dynamic exclusion function to avoid repeat scans of target ions and thereby shorten the analysis time of the samples. Another important function of dynamic exclusion was to obtain the fragment information of the co-eluting small peaks, which allowed compounds with low ionized potentials to be detected (Pan et al., 2015). The total ion current chromatogram of the methanol extract spiked with standards from *D. officinale* is presented, as obtained in the positive-ion mode under the optimized mass parameters and analytical conditions (**Figure S1a**). Generally, [M+H]^+^ or [M]^+^ was achieved through the soft ionization in the positive mode. The molecular-ion peaks of sarpagine and reserpine were separately extracted at *m/z* 311.1738 and 609.2772. The retention time of sarpagine and reserpine were at 12.95 min and 25.41 min, respectively (**Figure S1b** and **S1c**). The results indicated that the established method for the mass spectrometry was suitable for the separation and characterization of alkaloids.

Using a data-dependent MS^n^ scan method, the MS spectra of two standards were obtained from the Orbitrap analyzer, and the MS^n^ spectra were obtained from the LTQ. It was indicated that the π bonds on the indole groups of sarpagine and reserpine were easily dissociated to produce [M+H]^+^ peaks and produced massive hydrogen rearrangement. The ionization and electron transfer were observed on the position of the group cleavage because of the induced dissociation effect of the high-energy collisions. The putative fragmentation modes of sarpagine and reserpine are presented in Figure S2 and S3. The characteristic fragments with *m/z* 146 and 174 from the indole group were produced by the cleavage of sarpagine and reserpine, respectively. The characteristic ion *m/z* 195 cleaved on the ester bond was obtained to produce the 3,4,5-trimethoxybenzaldehyde group from the aglycone. Finally, forty-nine nitrogen-containing compounds, including seven alkaloids, were separated and tentatively identified from *D. officinale* (**Table S1**).

### Identification of the alkaloids

The small molecular compounds previously identified from *D. officinale* mainly include stilbenoids (bibenzyl and phenanthrene), phenols, lignans, nucleotides, lactones, flavonoids, steroids, and fatty acids. Based on the high resolution and sensitivity of LTQ-Orbitrap for the trace substances, coupled with the high selectivity of MCX for the medium-strength alkaline compounds, seven alkaloids identified from the methanol extract of *D. officinale* were divided into five classes based on the chemical structure, including tropane alkaloids, tetrahydroisoquinoline alkaloids, quinolizidine alkaloids, piperidine alkaloids, and spermidine alkaloids. The special fragmentation modes of different alkaloids in the ESI source were discussed below.

The tropane alkaloids consist of pyridine conjugated with piperidine. The experiment involved in DART-MS in-source CID at 90 V indicated that the characteristic ions of tropine were *m/z* 158, 142, 124, 93, and 67 (Lesiak et al., 2015). 2,3-Dihydroxynortropane was a kind of nortropane-type alkaloid that lacks an N-methyl group (Asano et al., 2001). Notably, the hydroxyl groups were present on its C-2 and C-3 in both the α- and β-configurations, as confirmed by a benzoate chirality method (Kusano et al., 2002). In this study, a strong [M+H-·OH]^+^ ion peak was observed under the set CID conditions. In addition to the characteristic ions *m/z* 93 and 67,and the stronger fragments *m/z* 113, 98 and 82 were observed (**Figure 2a**). The MS^2^ and MS^3^ spectra of 2,3-dihydroxynortropane were displayed, and [M+H-·OH]^+^ ion also appeared (**Figure 2b** and **2c**). However, the base peak of 2,3-dihydroxy-nortropane was the 2-methylpyrrole ion, which was further fragmented to get *m/z* 56. Finally, the compound was identified as 2,3-dihydroxynortropane or its isomers.Two tetrahydroisoquinoline alkaloids were identified as alamaridine and 1,2-dihydro-*O*-methyltazettine. The characteristic ions of tetrahydroisoquinoline group in alamaridine were *m/z* 177, 176, and 162 in the EI source, and the methyl group on C-5 was easier to lost compared to that on C-10. The MS^2^ and MS^3^ spectra of alamaridine are displayed respectively (**Figure 3a** and **3b**). These spectra indicated that the methyl groups on C-5 and C-10 were lost to produce the base peaks [M+H-2CH_3_-3H]^+^, which further lacked a hydroxyl group to generate *m/z* 248 in the MS^3^ spectrum. The C_2_H_3_ and ·OH groups were successively cleaved to generate *m/z* 221 and 204 on ring B and *m/z* 176 and 161 on ring C. 1,2-Dihydro-*O*-methyltazettine, also known as ungvedine, is a tazettine-type alkaloid (Kadyrov et al., 1979). The main fragments *m/z* 207 and 142 were generated by 1,2-dihydro-*O*-methyltazettine with equivalent intensity (**Figure 3c**). The *m/z* 142 further lacked the methyl group and simultaneously formed two double bonds to generate the base peak *m/z* 124 (**Figure 3d**). Previous studies have shown that the fragmentation pattern of tazettine-type alkaloid and its C3H7N group was easily lost to produce the characteristic ion [M+H-57]^+^ (Zhang et al., 2009). In Fig. 3c, 1,2-dihydro-*O*-methyltazettine was fragmented by RDA cleavage at both C-6, N-5 and C-4b, C-6a. 1,2-Dihydro-*O*-methyltazettine was cleaved at C-4b, C-12a but not on C-3, C-4, and thus did not produce [M+H-C_3_H_7_N]^+^. There was a coordination bond between nitrogen and oxygen presented in the quinolizidine alkaloid methyllagerine *N*-oxide. It was indicated that this bond easily led to the generation of the fragment [M-O]^+^ in the EI source (Lee et al., 2011). The oxygen was easily lost as well to produce the fragment *m/z* 437 in the ESI source (**Figure 4a**). The characteristic ions *m/z* 308 and 292 were generated by the cleavage of C-11 and C-22. According to the different cleavage positions on ring B, the characteristic ions *m/z* 220 and 163 were also generated. Meefarnine B was a spermidine alkaloid containing a polyamine heterocyclic ring. These alkaloids are generally considered to be synthesized by putrescine, which is produced by the degradation of L-arginine or L-ornithine (da Silva et al., 2015). The MS^2^ and MS^3^ spectra of meefarnine B indicate that the NH_3_ group on N-5 was cleaved to produce the fragment *m/z* 421 under the collision dissociation effect (**Figure 4b** and **4c**). The fragment *m/z* 292 was generated by the cleavage of the side chain on N-9 and was further cleaved to produce *m/z* 218. Besides, the fragment *m/z* 275 was generated by the cleavage of the *m/z* 421 on N-1 and C-2. The fragment *m/z* 204 was generated by the cleavage of the *m/z* 275 on N-9 and C-10, which further produced the characteristic ion *m/z* 147. In this study, a piperidine alkaloid anapheline was tentatively identified from *D. officinale*. The retention time of anapheline was 27.88 min, and the molecular weight of the [M+H]^+^ was 225.1945, which implies that its molecular formula was C_13_H_25_ON_2_ (**Figure S4**). Anapheline easily lost H_2_O to produce the fragment *m/z* 207 and the characteristic ion piperidine *m/z* 100 during the collision-induced dissociation. The NH3 group was lost to produce the *m/z* 83 in the MS^2^ spectrum. Based on the MS^2^ and MS^3^ spectra, a presumed fragmentation pathway of anapheline was proposed (**Figure 5**).

**Figure. 2.**
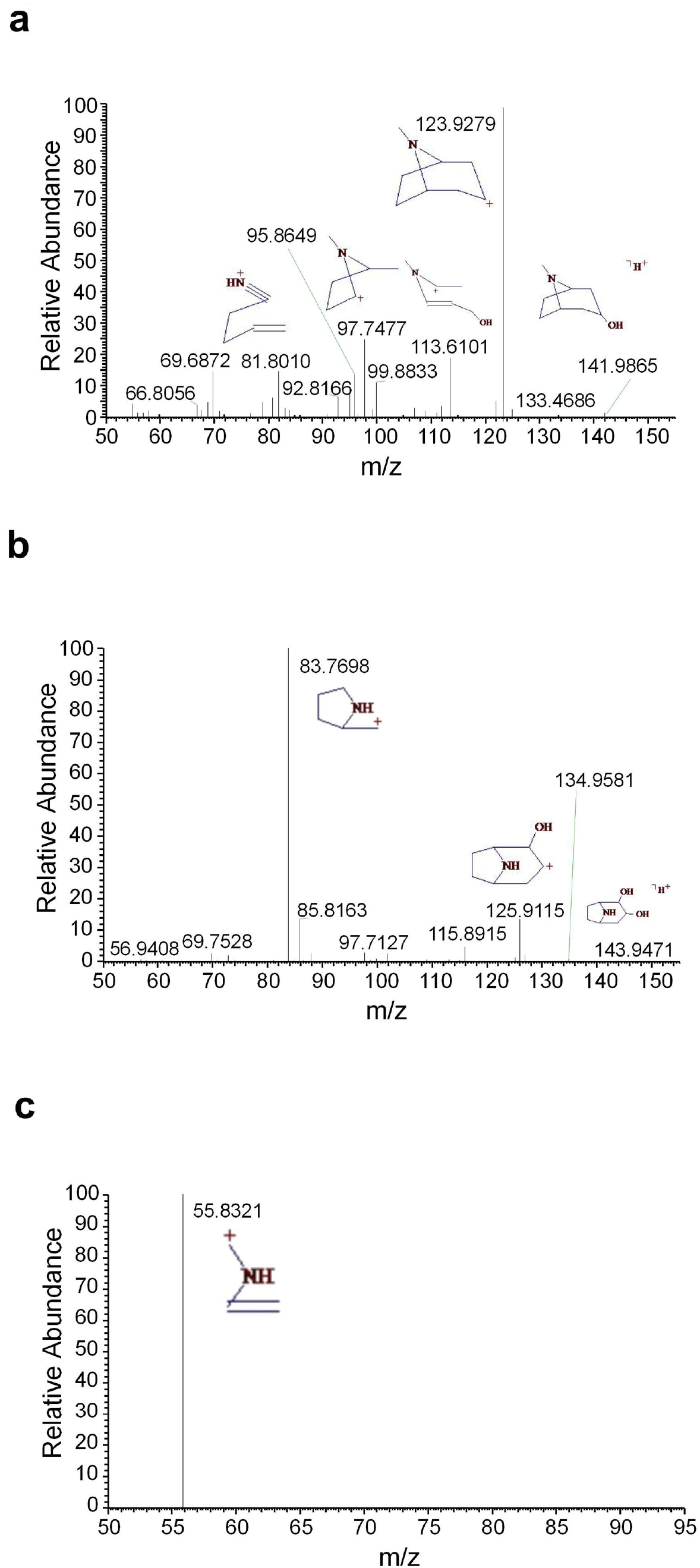
The mass spectra of tropine and 2,3-dihydroxynortropane. (a) The MS^2^ spectrum of tropine, (b) the MS^2^ spectrum of 2,3-dihydroxynortropane or an isomer, and (c) the MS^3^ spectrum of 2,3-dihydroxynortropane or an isomer.

**Figure. 3.**
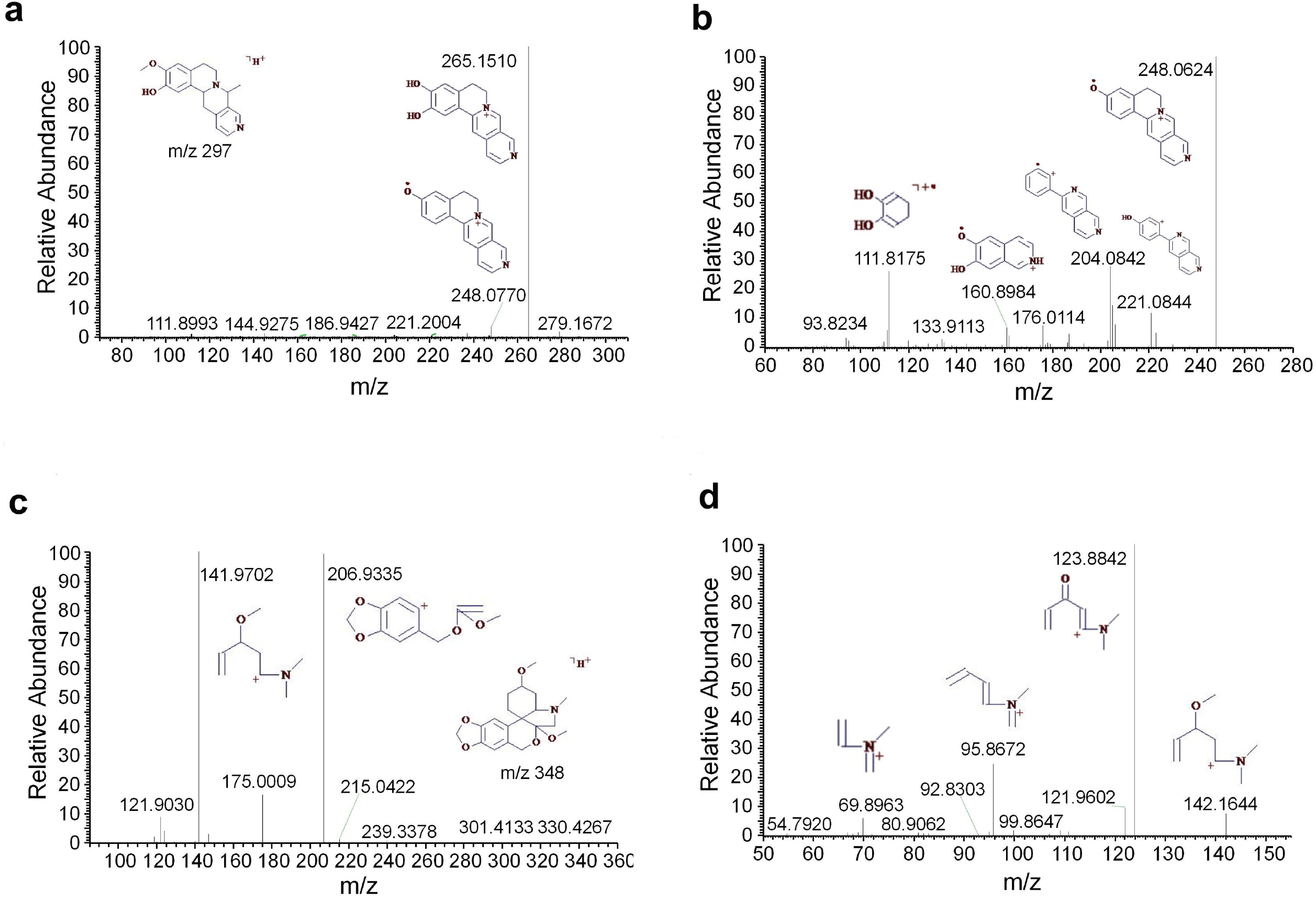
The mass spectra of alamaridine and 1,2-dihydro-*O*-methyltazettine. (a) The MS^2^ spectrum of alamaridine, (b) the MS^3^ spectrum of alamaridine, (c) the MS^2^ spectrum of 1,2-dihydro-*O*-methyltazettine, and (d) the MS^3^ spectrum of 1,2-dihydro-*O*-methyltazettine.

**Figure. 4.**
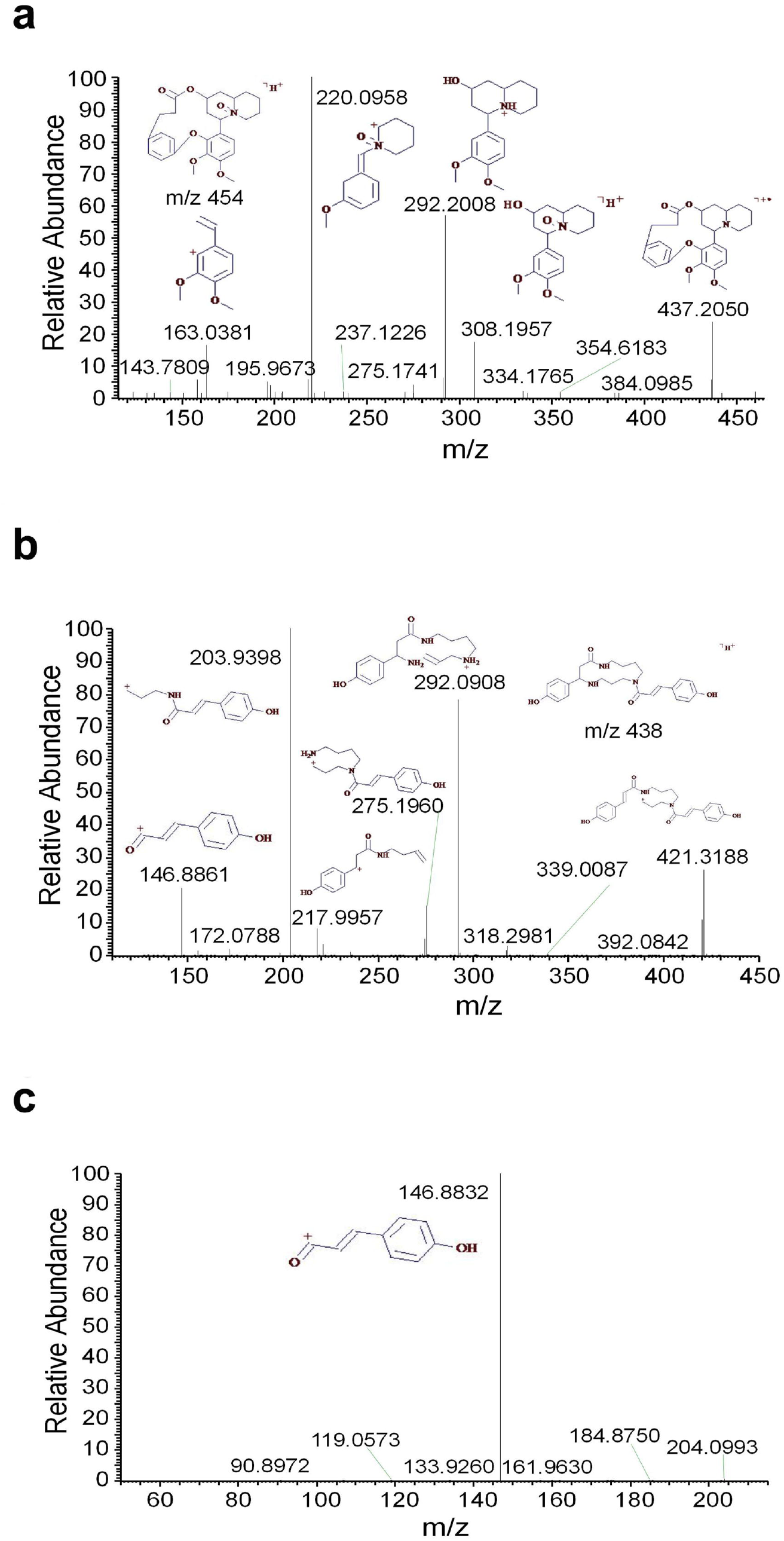
The mass spectra of methyllagerine *N*-oxide and meefarnine B. (a) The MS^2^ spectrum of methyllagerine *N*-oxide, (b) the MS^2^ spectrum of meefarnine B, and (c) the MS^3^ spectrum of meefarnine B.

**Figure. 5.**
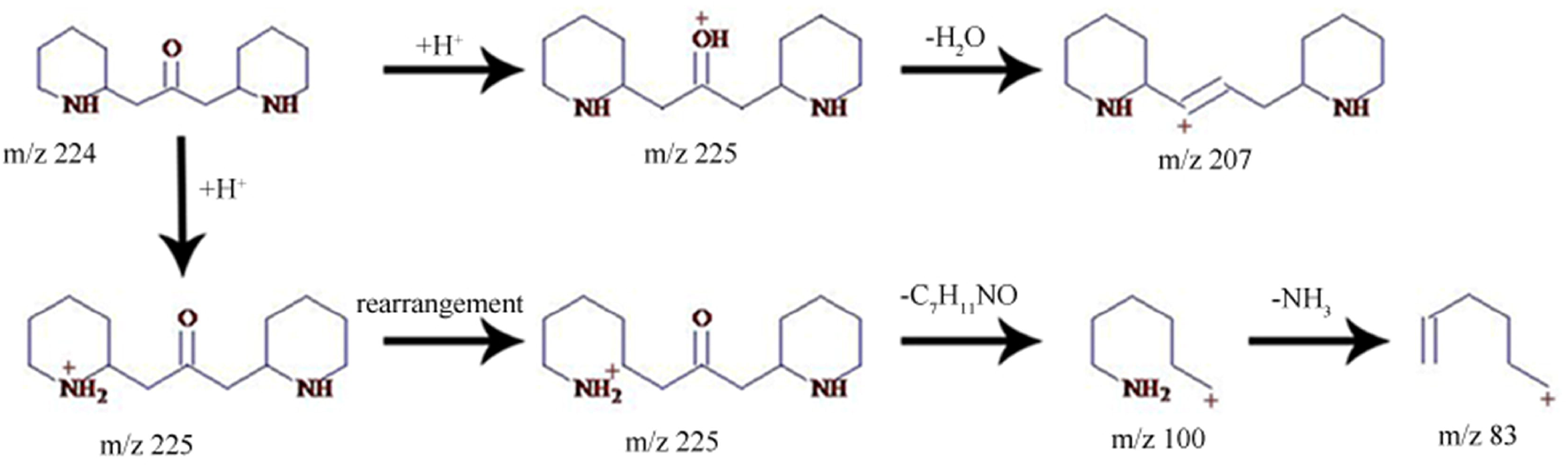
The putative fragmentation pathway of anapheline.

### Identification of the guanidines

Three guanidine compounds were separated and determined from the methanol extract of *D. officinale* for the first time. The MS^2^ and MS^3^ spectra of 4-guanidinobutanoic acid were displayed respectively (**Figure 6a** and **6b**). The mass spectra indicated that 4-guanidinobutanoic acid easily lost H_2_O to form the base peak *m/z* 128 and produced the characteristic ions *m/z* 104, 87 and 60. The *m/z* 128 was further cleaved to produce the *m/z* 111 and 86. These results were consistent with those reported by the previous study (Nikolić et al., 2012). Cyclocimipronidine is a cyclic guanidine compound separated from *Cimicifuga racemosa*. Some studies showed that the characteristic ions of cyclocimipronidine, a guanidine alkaloid in black cohosh, were the *m/z* 112, 95, 94, 70 and 67 (Gödecke et al., 2009). In addition to the aforementioned characteristic ions, cyclocimipronidine easily lost H_2_O to generate the fragments *m/z* 136 and 84 (**Figure 6c** and **6d**). The MS^2^ and MS^3^ spectra of *N*-*tert*-butyloxycarbonyl guanidine were displayed in Fig. 6e and 6f, respectively. It was indicated that the characteristic ions were the *m/z* 142, 125, 104 and 87.

**Figure. 6.**
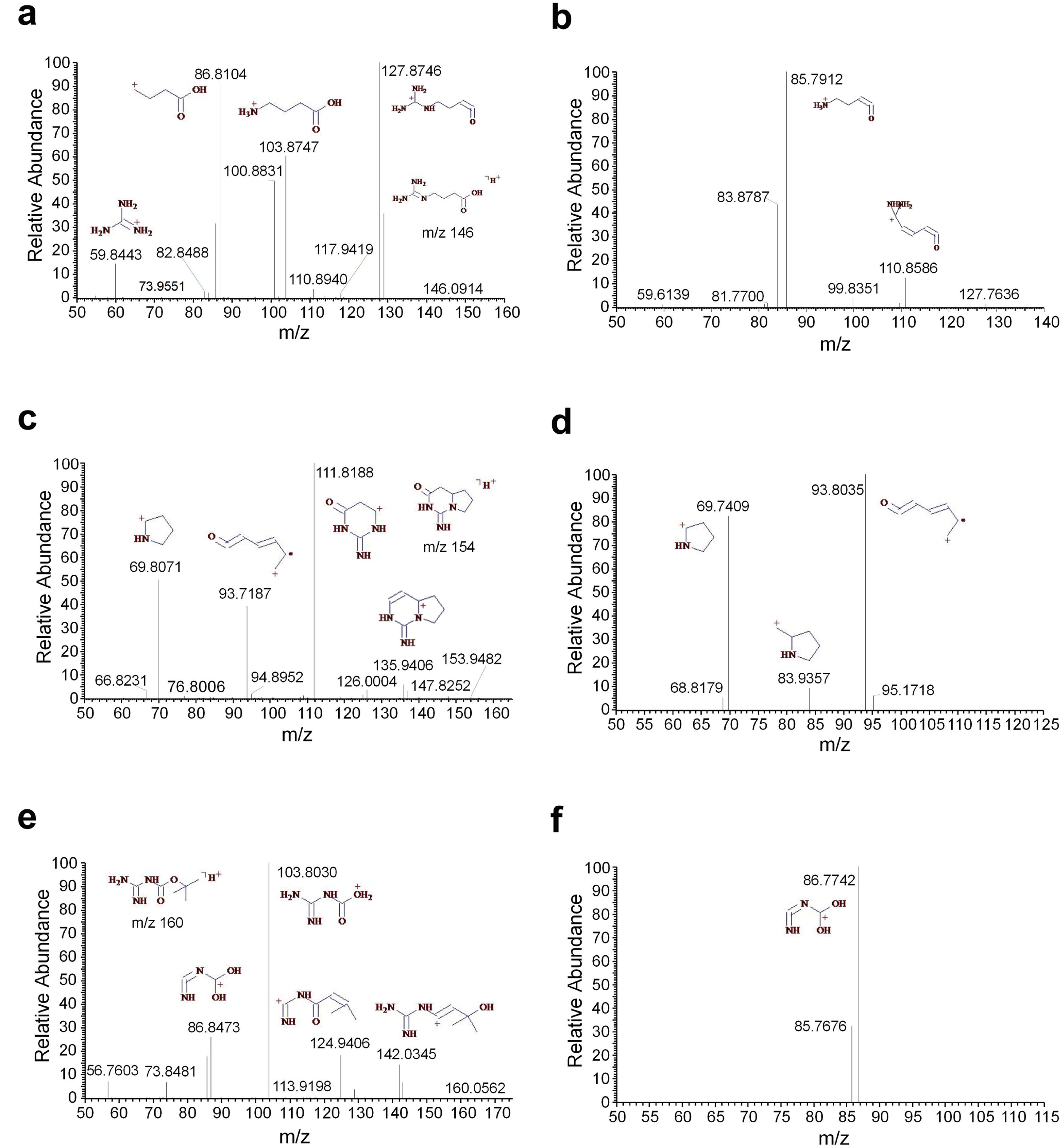
The mass spectra of the guanidines. (a) The MS^2^ spectrum of 4-guanidinobutanoic acid, (b) the MS^3^ spectrum of 4-guanidinobutanoic acid, (c) the MS^2^ spectrum of cyclocimipronidine, (d) the MS^3^ spectrum of cyclocimipronidine, (e) the MS^2^ spectrum of *N*-*tert*-butyloxycarbonyl guanidine, and (f) the MS^3^ spectrum of *N*-*tert*-butyloxycarbonyl guanidine.

### Identification of the nucleotides and derivatives

Nucleotides were composed of purine or pyrimidine linked with ribose. adenosine, uridine and guanosine had been separated from *D. officinale* in a previous study. In this study, five nucleotide derivatives were tentatively identified. 2-Hydroxyadenine was an adenine derivative that easily lost an NH_3_ group to produce the characteristic ions *m/z* 135 and 110 (**Figure S5a**). The fragments *m/z* 107, 92 and 77 were generated in the MS^3^ spectrum (**Figure S5b**). Besides 2-hydroxyadenine, another four nucleotide derivatives were identified in the study. Their MS^2^ and MS^3^ spectra indicated that the neutral loss of *m/z* 136 resulted from the loss of aglycones (**Figure S5 c-j**). Comparison with the multistage mass spectra of the METLIN database (https://metlin.scripps.edu/) resulted in the identification of adenosine, guanosine, 1-methyladenosine and cytidine.

### Identification of the sphingolipids

Sphingolipid is a structurally complex lipid containing a long-chain sphingosine skeleton, which forms the plasma and vacuole membranes of plant cells. Four phytosphingosines were separated and identified from *D. officinale*. Because sphingosine contained some hydroxyl groups, the characteristic ion [M+H-H_2_O]^+^ was observed during the dissociation (**Figure S6a-h**). By aligning against the Dictionary of Natural Products Database (http://dnp.chemnetbase.com), four sphingolipids, including 2-amino-1,3-hexadecanediol, 2-amino-3-tetradecanol, 2-amino-1,3,4-octadecanetriol and 2-amino-1,3,4-hexadecanetriol, were identified in the experiment.

### Identification of the amino acids and their derivatives

Eight amino acids—l-lysine, l-histidine, l-arginine, l-valine, l-isoleucine, l-tyrosine, l-phenylalanine and l-tryptophan—were identified from *D. officinale* based on accurate molecular weights and the multistage mass spectra of standards from the MassBank mass spectrum library (http://www.massbank.jp/QuickSearch.html) (**Table S1**). 4-Hydroxy-l-threonine and urocanic acid were derivatives of L-threonine and L-histidine that easily lost H_2_O to produce the base peaks *m/z* 118 and 121. The putative fragment ions generated during their cleavage were displayed in **Figure S7a** and **S7b**, respectively. In addition to the derivatives of l-threonine and l-histidine, four additional arginine derivatives were identified in this study. *N^2^*-Fructopyranosylarginine was an arginine monoglycoside and the byproduct generated by a Maillard reaction between reducing sugars (fructose or glucose) and arginine upon heating (Suzuki et al., 2004). Because fructose contained several hydroxyl groups, H_2_O was easily lost to produce the characteristic ions *m/z* 319, 301 and 283. The fragments *m/z* 257 and 239 were produced by the cleavage of the guanidino group on the glycoside. N^2^-Fructopyranosylarginine was also cleaved to generate the characteristic ion *m/z* 175 on the N-glucosidic band and further produced the aglycone *m/z* 158 via the neutral loss of fructose (**Figure S7c-d**). The NH_3_ group of *N*-methylarginine was lost to produce the base peak *m/z* 172, as a result of the dissociation. According to the different cleavages on the R group, the characteristic ions *m/z* 158, 133, 116 and 74 were generated (**Figure S7e**). The characteristic ions *m/z* 126, 115, 97 and 70 were also generated in the MS^3^ spectrum by the cleavage of the *m/z* 172 fragment (**Figure S7f**). The ·OH group of *N^α^*-acetylarginine was lost to produce the characteristic ion *m/z* 200. The fragments *m/z* 175 and 157 were generated by the different cleavages on the R group. The characteristic ions *m/z* 139, 115 and 70 were produced by the cleavage of amino and carboxyl groups linked with the chiral carbons. The ·OH group of the *m/z* 200 was lost to produce the *m/z* 183 in the MS^3^ spectrum, and the other fragments were similar to those in the MS^2^ spectrum (**Figure S7g-h**). *N^2^*-(3-Hydroxysuccinoyl)arginine was formed by the substitution of the 3-hydroxysuccinoyl group on the N-terminal of arginine. The typical feature of its cleavage was the loss of 3-hydroxysuccinoyl group to produce the characteristic ion *m/z* 175 (**Figure S7i**), which was further cleaved to generate the fragments *m/z* 158, 139 and 60 (**Figure S7j**).

### Identification of some isomers

The accurate molecular weights of the compounds were provided by high-resolution mass spectrometry, which allowed us to deduce the speculative molecular formulas. However, the identification of isomers was less effective. Two pairs of isomers were identified in this study based on their different retention times and distinct multistage fragments. The formula of both L-homophenylalanine and β-phenyl-γ-aminobutyric acid was C_10_H_13_O_2_N, but their retention times were at 6.79 and 12.47 min, respectively (**Table S1**). The MS^2^ spectra of L-homophenylalanine and β-phenyl-γ-aminobutyric acid are displayed in **Figure S8a** and **S8c**, respectively. These mass spectra indicated that the NH3 groups were easily lost to form the characteristic ion *m/z* 163. However, the fragment *m/z* 120 produced in the spectrum of β-phenyl-γ-aminobutyric acid was not observed in that of L-homophenylalanine. The MS^3^ spectra presented different fragments, which indicated two different compounds (**Figure S8b** and **S8d**). The formula of both carbetamide and *N^δ^*-benzoylornithine were C_12_H_16_O_3_N_2_, but their retention times were at 9.77 and 14.93 min, respectively (**Table S1**). The MS^2^ and MS^3^ spectra of carbetamide were displayed in **Figure S8e** and **S8f**, respectively. The MS^2^ and MS^3^ spectra of *N^δ^*-benzoylornithine were displayed in Fig. S8g and S8h, respectively. These two isomers exhibited different fragmentation patterns. The characteristic ions of carbetamide were the *m/z* 120, 103 and 93 ions, whereas the characteristic ions of *N^δ^*-benzoylornithine were the *m/z* 219, 205, 175, 158 and 148 ions. The isomerism of the dipeptides resulted from the chirality of the α-carbon and the different positions of peptide bonds between two peptides. In the study, eight dipeptides were tentatively identified. Most of these dipeptides had more than one retention time, suggesting that the isomers existed.

### Identification of the nitrogen-containing glycosides

By comparing our spectra with the Metlin and Dictionary of Natural Products databases, we tentatively identified four nitrogen-containing glycosides from *D. officinale*. The neutral loss of H_2_O in 5’-*O*-β-d-glucosylpyridoxine was easy to produce the characteristic ion *m/z* 314. The fragment *m/z* 171 was generated by the cleavage on the O-glucosidic bond and further produced the aglycone *m/z* 152 by the neutral loss of glucose (**Figure S9a**). The H_2_O on the aglycone of 4-*O*-β-d-glucopyranoside-2,4-benzoxazolediol was easily lost to produce the base peak *m/z* 296. The benzoxazole ion *m/z* 136 was generated by the cleavage on the O-glucosidic bond (**Figure S9b**). The *m/z* 296 was cleaved to generate the characteristic ions *m/z* 119 and 94 (**Figure S9c**). The MS^2^ and MS^3^ spectra of *O*-(tri-*O*-acetyl-α-l-rhamnopyranoside)- (4-hydroxybenzyl) methylcarbamic acid were displayed in **Figure S9d** and **S9e**, respectively. These mass spectra indicated that the characteristic ion *m/z* 322 was generated by the cleavage of the ester bond and the side chain. The *m/z* 322 was cleaved to produce the fragments *m/z* 304 and 160 in the MS^3^ spectrum. Passicapsin was composed of one glucose and a nitrogen-containing aglycone that easily produced the aglycone *m/z* 238 via the neutral loss of glucose. The characteristic ions *m/z* 286 and 148 were cleaved at the O-glucosidic bond between the side chain and the dideoxy glucoside (**Figure S9f**). The fragments *m/z* 148 and 106 in the MS^3^ spectrum were produced by the cleavage of the aglycone *m/z* 238 on the O-glucosidic bond. The H_2_O of the aglycone *m/z* 238 was lost to produce the fragments *m/z* 220 and 212, and the dideoxy glucoside group was further cleaved to generate the *m/z* 190 and 174 (**Figure S9g**).

## Conclusions

The alkaloids and precursors of *D. officinale* with MeJA treatment were primarily studied by the SPE-HPLC–LTQ-Orbitrap technique. The extraction and purification of the target compounds was optimized, and the suitable conditions of ESI-CID-MS^n^ for the mass spectrometric detection were established. Based on the accurate mass weight with MS^n^ fragmentation information provided by the LTQ-Orbitrap mass spectrometer, forty-nine compounds, including amino acids derivatives, dipeptides, nucleotides, guanidines, alkaloids, sphingolipids and several nitrogen-containing glycosides, were tentatively identified from *D. officinale*. Seven alkaloids were identified from *D. officinale*. The combination of alkaloids and their precursors were associated to construct a metabolic network for the interpretation of biosynthetic pathways. These findings will serve as a technical guidance for the analyses and drug discoveries of *Dendrobium* alkaloids.

## Acknowledgement

We are grateful to Mrs. Airong Feng of University of Science and Technology of China for her support of the work on mass spectra interpretation.

## Author contributions

CS and GHL discussed the writing plan. CS and YPZ drafted the manuscript. CS and MAM edited the manuscript. GHL and CS acquired the funding. All authors have read, reviewed, and approved the submitted version.

**Figure. S1**. The mass spectrum of *D. officinale* extract with the standards. (a) The total ion current chromatogram of the methanol extract from *D. officinale* spiked with standards, (b) the extracted ion current chromatogram of *m/z* 311.1738, and (c) the extracted ion current chromatogram of *m/z* 609.2772.

**Figure. S2**. The putative fragmentation pathway of sarpagine. π denotes π-bond dissociation; i denotes inductive cleavage; rH_R_ denotes charge-remote rearrangement; rH_B_ denotes α,β-charge-site rearrangement; rH_C_ denotes γ-charge-site rearrangement; and rH_1,2_ denotes radical-site rearrangement.

**Figure. S3**. The putative fragmentation pathway of reserpine. π denotes π-bond dissociation; i denotes inductive cleavage; rH_R_ denotes charge-remote rearrangement; and rH_C_ denotes γ-charge-site rearrangement.

**Figure. S4**. The chromatogram and mass spectra of anapheline. (a) The extracted ion chromatogram of anapheline, (b) the MS spectrum of anapheline, (c) the MS^2^ spectrum of anapheline, and (d) the MS^3^ spectrum of anapheline.

**Figure. S5**. The mass spectra of nucleotides and derivatives. (a) The MS^2^ spectrum of 2-hydroxyadenine, (b) the MS^3^ spectrum of 2-hydroxyadenine, (c) the MS^2^ spectrum of adenosine, (d) the MS^3^ spectrum of adenosine, (e) the MS^2^ spectrum of guanosine, (f) the MS^3^ spectrum of guanosine, (g) the MS^2^ spectrum of 1-methyladenosine, (h) the MS^3^ spectrum of 1-methyladenosine, (i) the MS^2^ spectrum of cytidine, and (j) the MS^3^ spectrum of cytidine.

**Figure. S6**. The mass spectra of sphingolipids. (a) The MS^2^ spectrum of 2-amino-1,3-hexadecanediol, (b) the MS^3^ spectrum of 2-amino-1,3-hexadecanediol, (c) the MS^2^ spectrum of 2-amino-3-tetradecanol, (d) the MS^3^ spectrum of 2-amino-3-tetradecanol, (e) the MS^2^ spectrum of 2-amino-1,3,4-octadecanetriol, (f) the MS^3^ spectrum of 2-amino-1,3,4-octadecanetriol, (g) the MS^2^ spectrum of 2-amino-1,3,4-hexadecanetriol, and (h) the MS^3^ spectrum of 2-amino-1,3,4-hexadecanetriol.

**Figure. S7**. The mass spectra of amino acids and their derivatives. (a) The MS^2^ spectrum of 4-hydroxy-l-threonine, (b) the MS^2^ spectrum of urocanic acid, (c) the MS^2^ spectrum of *N^2^*-fructopyranosylarginine, (d) the MS^3^ spectrum of *N^2^*-fructopyranosylarginine, (e) the MS^2^ spectrum of *N*-methylarginine, (f) the MS^3^ spectrum of *N*-methylarginine, (g) the MS^2^ spectrum of *N^α^*-acetylarginine, (h) the MS^3^ spectrum of *N^α^*-acetylarginine, (i) the MS^2^ spectrum of *N^2^*-(3-hydroxysuccinoyl)arginine, and (j) the MS^3^ spectrum of *N^2^*-(3-hydroxysuccinoyl)arginine.

**Figure. S8**. The comparative mass spectrum of some isomers. (a) The MS^2^ spectrum of L-homophenylalanine, (b) the MS^3^ spectrum of L-homophenylalanine, (c) the MS^2^ spectrum of β-phenyl-γ-aminobutyric acid, (d) the MS^3^ spectrum of β-phenyl-γ-aminobutyric acid, (e) the MS^2^ spectrum of carbetamide, (f) the MS^3^ spectrum of carbetamide, (g) the MS^2^ spectrum of *N^δ^*-benzoylornithine, and (h) the MS^3^ spectrum of *N^δ^*-benzoylornithine.

**Figure. S9**. The mass spectra of nitrogen-containing glycosides. (a) The MS^2^ spectrum of 5’-*O-β*-d-glucosylpyridoxine, (b) the MS^2^ spectrum of 4-*O*-*β*-d-glucopyranoside-2,4-benzoxazolediol, (c) the MS^3^ spectrum of 4-*O*-*β*-d-glucopyranoside-2,4-benzoxazolediol, (d) the MS^2^ spectrum of *O*-(tri-*O*-acetyl-*α*-l-rhamnopyranoside)-(4-hydroxy benzyl)methylcarbamic acid, (e) the MS^3^ spectrum of *O*-(tri-*O*-acetyl-*α*-l-rhamnopyranoside)-(4-hydroxy benzyl)methylcarbamic acid, (f) the MS^2^ spectrum of passicapsin, and (g) the MS^3^ spectrum of passicapsin.

**Table S1.** Identification of the main chemical compounds from *D.officinale*

