## Supplementary figures and images for "Identification of Alkaloids and Related Intermediates of *Dendrobium officinale* by Solid-Phase Extraction Coupled with High-Performance Liquid Chromatography Tandem Mass Spectrometry"

### Supplemental Figure 1

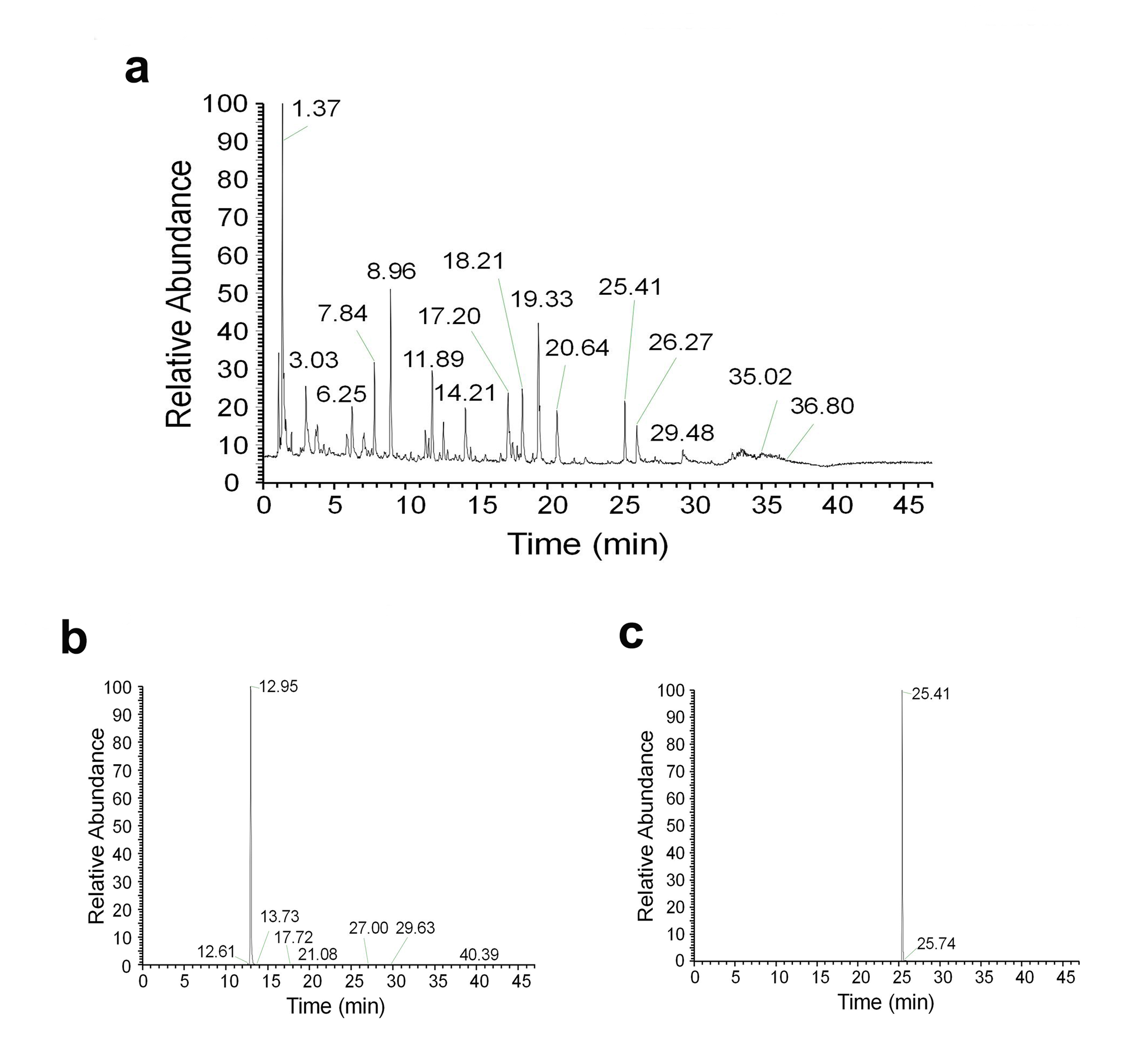

### Supplemental Figure 2

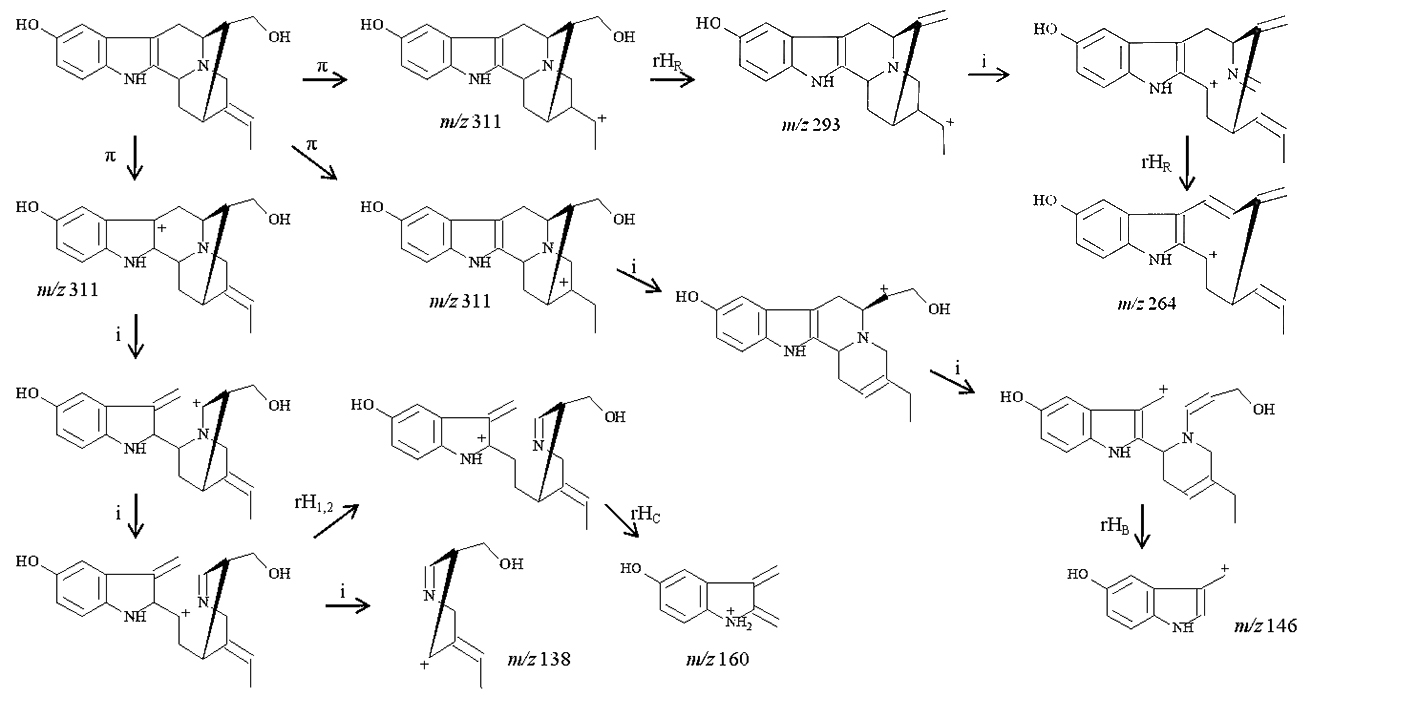

### Supplemental Figure 3

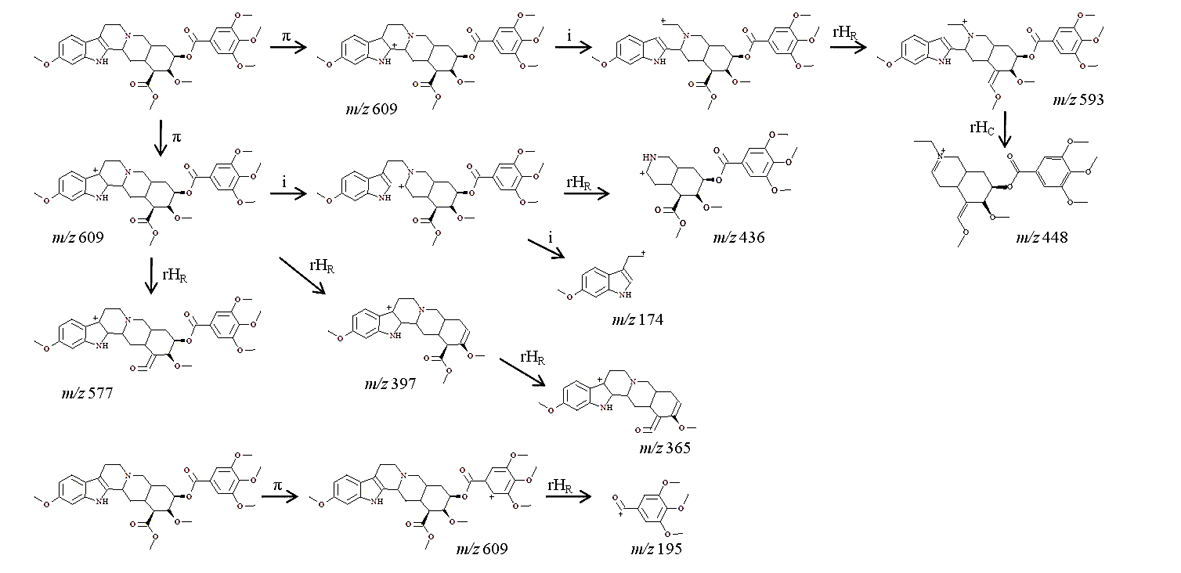

### Supplemental Figure 4

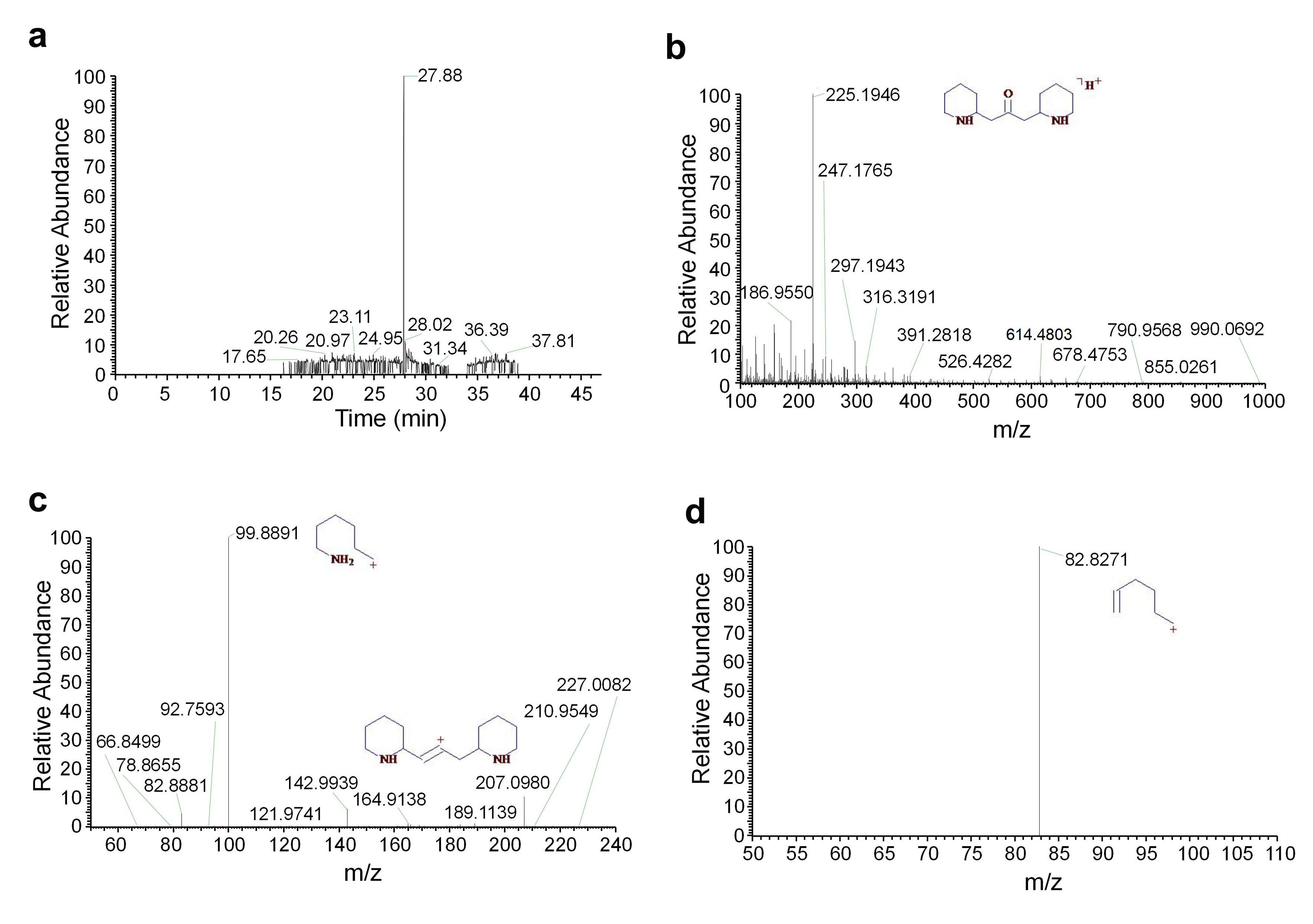

### Supplemental Figure 5

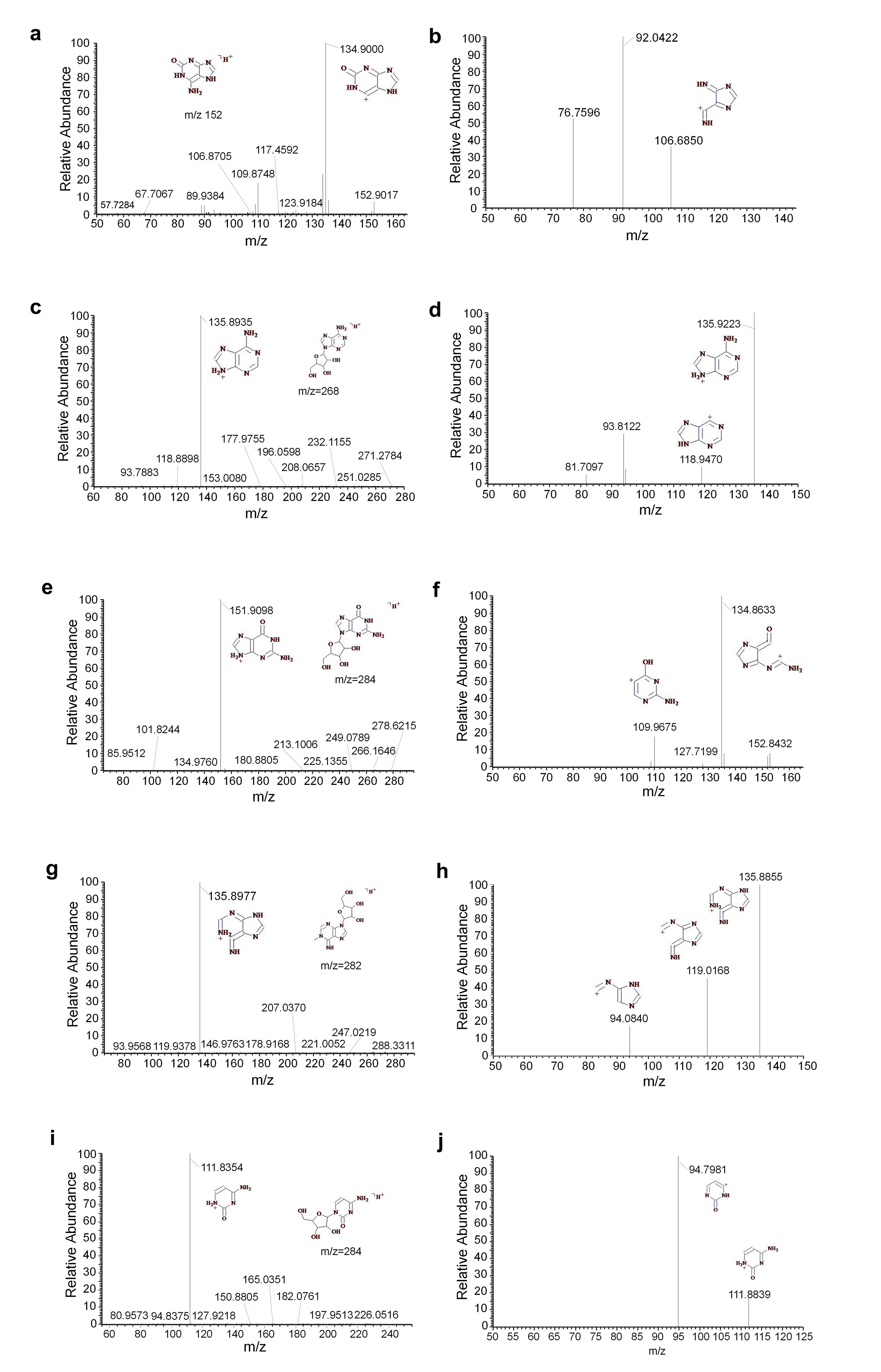

### Supplemental Figure 6

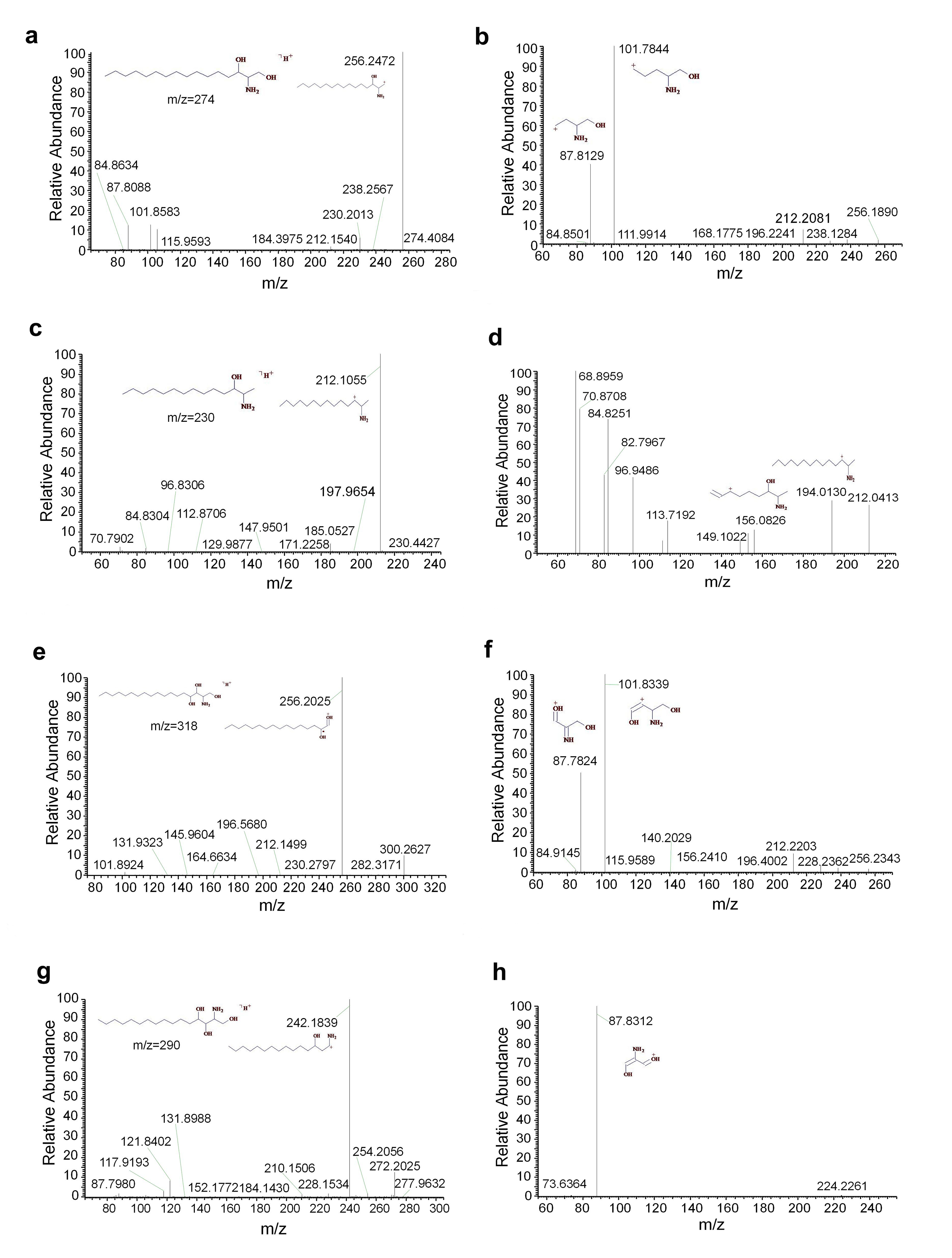

### Supplemental Figure 7

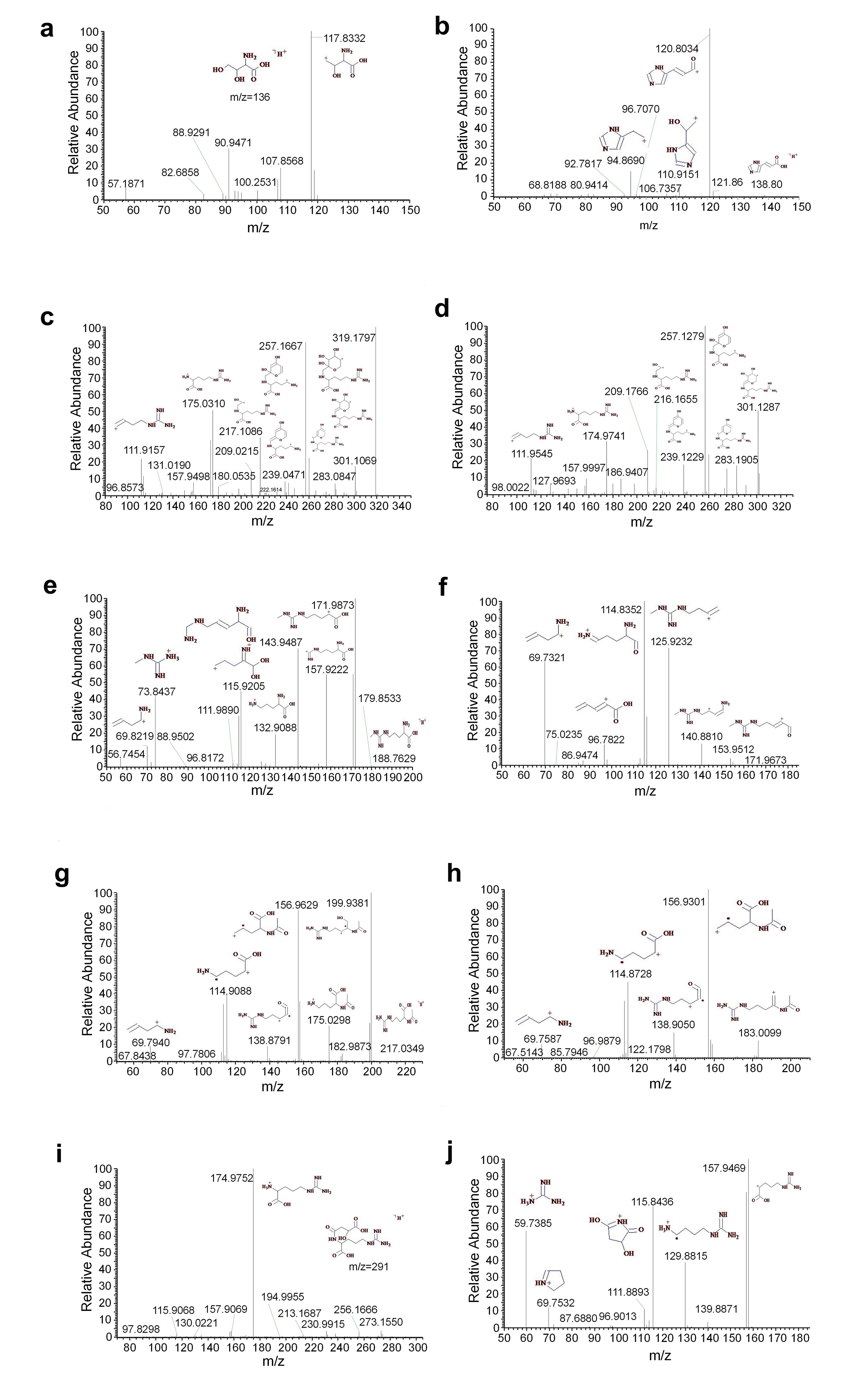

### Supplemental Figure 8

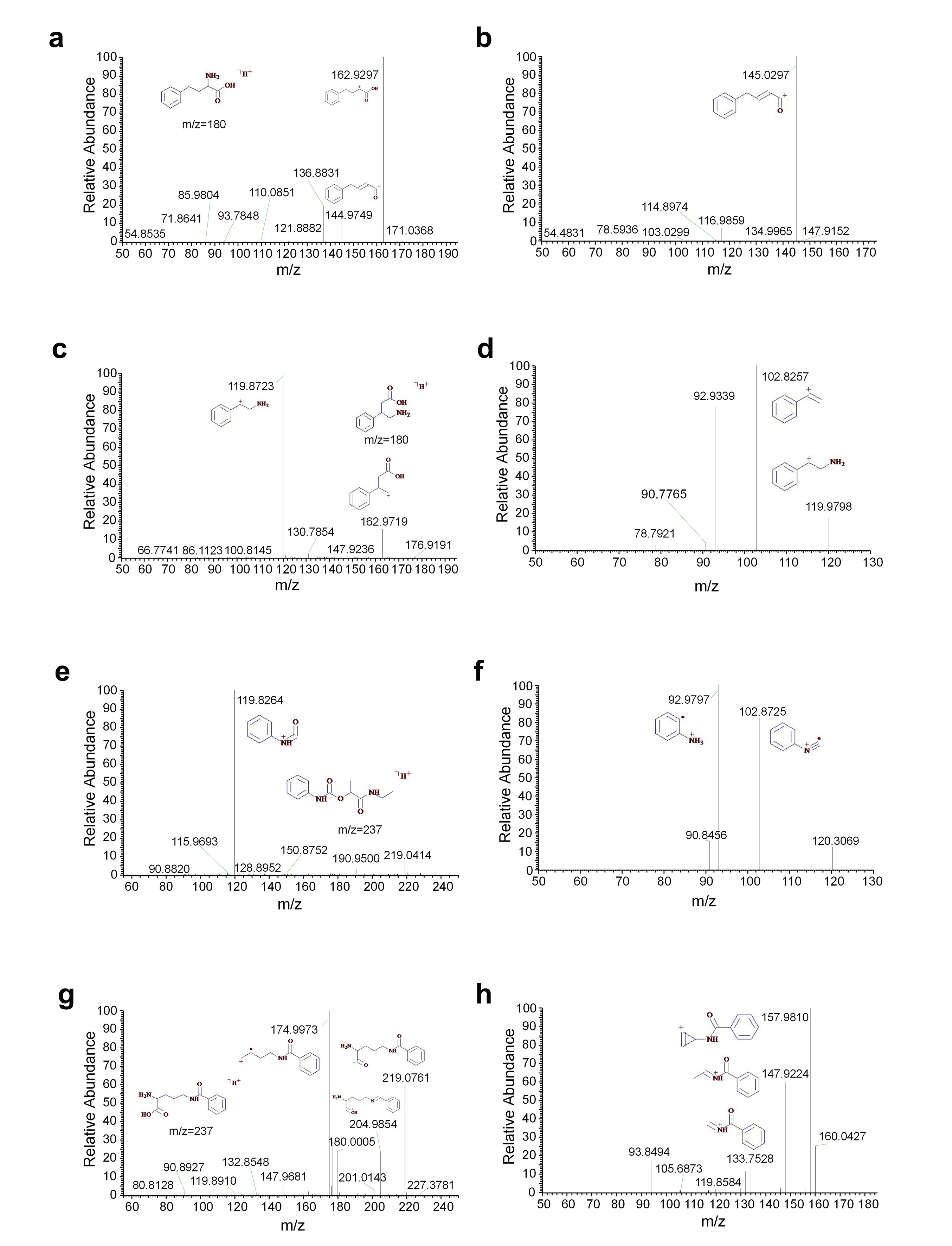

### Supplemental Figure 9

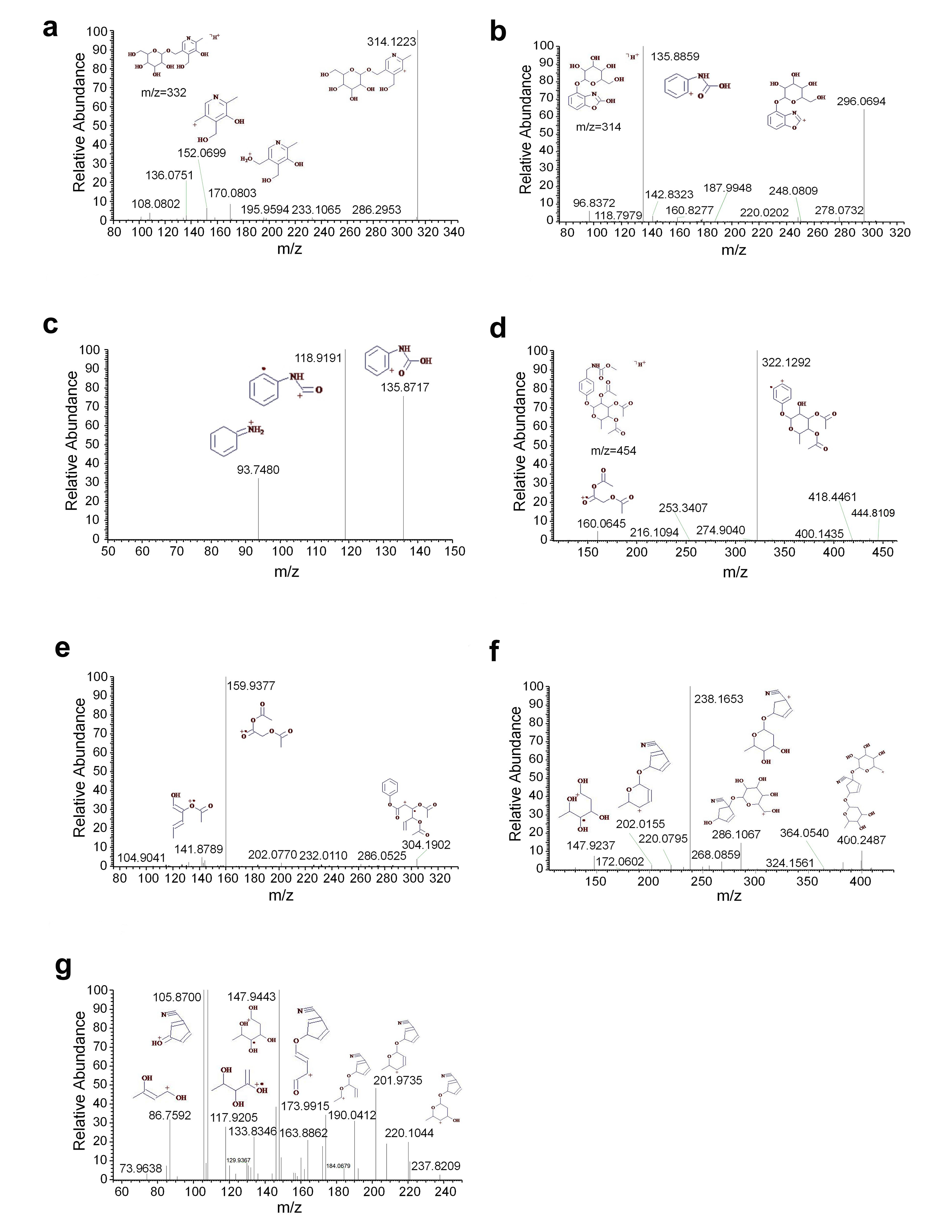
