## Supplemental Table 1 for "Identification of Alkaloids and Related Intermediates of *Dendrobium officinale* by Solid-Phase Extraction Coupled with High-Performance Liquid Chromatography Tandem Mass Spectrometry"

**Table S1. Identification of main chemical compounds from *D.officinale***

| ***t*_R_ (min)** | **[M+H]^+^** | **Formula** | **Error (mmu)** | **Fragment ions** | **Identification** |
| --- | --- | --- | --- | --- | --- |
| 2.64/3.19 | 118.0855 | C_5_H_12_O_2_N | -0.795 | MS^2^[118]:72 MS^3^[118→72]:55 | l-Valine |
| 5.89/6.24 | 132.1010 | C_6_H_14_O_2_N | -0.905 | MS^2^[132]:86 MS^3^[132→86]:69;58 | l-Isoleucine |
| 3.81/7.73 | 136.0609 | C_4_H_10_O_4_N | -0.832 | MS^2^[136]:119;118;108;107;100;91;57 | 4-Hydroxy- l-threonine |
| 5.00/6.42 | 139.0494 | C_6_H_7_O_2_N_2_ | -0.784 | MS^2^[139]: 121;111;95 | Urocanic acid |
| 2.81/3.02 | 142.1218 | C_8_H_16_ON | -0.841 | MS^2^[142]: 124;114;98;82;70;55 | Tropine or its isomers |
| 5.14/5.65 | 144.1011 | C_7_H_14_O_2_N | -0.835 | MS^2^[144]:126;86;84  MS^3^[144→84]:56 | 2,3-Dihydroxynortropane or its isomers |
| 3.71 | 146.0914 | C_5_H_12_O_2_N_3_ | -0.973 | MS^2^[146]: 128;104;101;87;86;60 MS^3^[146→128]:111;86;84 | 4-Guanidinobutanoic acid |
| 1.21 | 147.1120 | C_6_H_15_O_2_N_2_ | -0.854 | MS^2^[147]:130; 129;84 MS^3^[147→130]:84 | l-Lysine |
| 4.06 | 152.0559 | C_5_H_6_ON_5_ | -0.806 | MS^2^[152]:136;135;134;124;110;109;90  MS^3^[152→135]:107;92;77 | 2-Hydroxyadenine |
| 5.56 | 154.0967 | C_7_H_12_ON_3_ | -0.839 | MS^2^[154]:136;112;94;70 MS^3^[154→112]:95;94;84;70;69;67 | Cyclocimipronidine |
| 1.29 | 156.0759 | C_6_H_10_O_2_N_3_ | -0.873 | MS^2^[156]: 110; 95 MS^3^[156→110]: 93; 83 | l-Histidine |
| 6.42 | 160.1070 | C_6_H_14_O_2_N_3_ | -1.023 | MS^2^[160]:143;142;125;104;87;86;74 MS^3^[160→104]:87;86 | *N-tert*-Butyloxycarbonyl guanidine |
| 8.88 | 166.0853 | C_9_H_12_O_2_N | -1.005 | MS^2^[166]:120 MS^3^[166→120]:120;103;93;91 | l-Phenylalanine |
| 1.36 | 175.1178 | C_6_H_15_O_2_N_4_ | -1.182 | MS^2^[175]: 158;140;130;116;112;70;60 | l-Arginine |
| 12.47 | 180.1007 | C_10_H_14_O_2_N | -1.225 | MS^2^[180]:163;120 MS^3^[180→120]:120;103;93 | β-Phenyl-γ-aminobutyric acid |
| 6.79 | 180.1008 | C_10_H_14_O_2_N | -1.075 | MS^2^[180]:163;145;137 MS^3^[180→163]:145;117 | l-Homophenylalanine |
| 7.00 | 182.0801 | C_9_H_12_O_3_N | -1.06 | MS^2^[182]:165;147;136 | l-Tyrosine |
| 1.60 | 189.1336 | C_7_H_17_O_2_N_4_ | -0.982 | MS^2^[189]:172;171;158;144;133;116;115;74;70  MS^3^[189→172]:141;126;116;115 | *N^g^*-Methyl-l-arginine |
| 7.34/7.65/8.73/9.43 | 203.1379 | C_9_H_19_O_3_N_2_ | -1.199 | MS^2^[203]:185;157;132;86 MS^3^[203→132]:86;69 | Alanyl-isoleucine or its isomers |
| 11.86 | 205.0960 | C_11_H_13_O_2_N_2_ | -1.144 | MS^2^[205]:188 MS^3^[205→188]:170;146;144 | l-Tryptophan |
| 4.09 | 217.1285 | C_8_H_17_O_3_N_4_ | -1.037 | MS^2^[217]:200;199;175;158;157;139;115;113;70 MS^3^[217→200]:183;158;157;139;115; 113;70 | *N^α^*-Acetyl-l-arginine |
| 8.04 | 217.1534 | C_10_H_21_O_3_N_2_ | -1.239 | MS^2^[217]:199;171;118;72 | Valyl-valine |
| 27.88 | 225.1945 | C_13_H_25_ON_2_ | -1.59 | MS^2^[225]:207;143;100;83; MS^3^[225→100]:83 | Anapheline |
| 26.46 | 230.2462 | C_14_H_32_ON | -1.641 | MS^2^ [230]:212;185;  MS^3^[230→212]: 194; 169;156;125;111;97;85;71;69 | 2-Amino-3-tetradecanol |
| 9.98/10.6/11.48 | 231.1690 | C_11_H_23_O_3_N_2_ | -1.309 | MS^2^[231]:213;185;132;72 MS^3^[231→72]:55 | Valyl-leucine or its isomers |
| 14.93 | 237.1219 | C_12_H_17_O_3_N_2_ | -1.469 | MS^2^[237]:219;205;180;177;175;148  MS^3^[237→175]:160;158;148;134;132;94 | *N^δ^*-Benzoylornithine |
| 9.77 | 237.1219 | C_12_H_17_O_3_N_2_ | -1.469 | MS^2^[237]:219;120 MS^3^[237→120]: 120;103;93;91 | Carbetamide |
| 4.25 | 244.0913 | C_9_H_14_O_5_N_3_ | -1.487 | MS^2^[244]:112 MS^3^[244→112]:112;95 | Cytidine |
| 12.55/13.24/13.59/14.35 | 245.1844 | C_12_H_25_O_3_N_2_ | -1.599 | MS^2^[245]:227;199;132;86  MS^3^[245→86]:69 | Leucyl-isoleucine or its isomers |
| 10.37 | 253.1280 | C_11_H_17_O_3_N_4_ | -1.557 | MS^2^[235]:236;235;225;208;193;192;191;181;165;164;147;112 MS^3^[235→236]:218;209;193;192;191;175;165; 147;121;70 | Prolyl-histidine |
| 11.41/12.73 | 265.1531 | C_14_H_21_O_3_N_2_ | -1.529 | MS^2^[265]:248;206;177;152;114;89 MS^3^[265→248]:177;145 | Phenylalanyl-valine or its isomers |
| 7.73 | 268.1025 | C_10_H_14_O_4_N_5_ | -1.58 | MS^2^[268]:136 MS^3^[268→136]:136;119;94;82 | Adenosine |
| 26.34 | 274.2720 | C_16_H_36_O_2_N | -2.026 | MS^2^[274]: 256;230;102;88;  MS^3^[274→256]: 238;212;102;88 | 2-Amino-1,3-hexadecanediol |
| 14.93/15.67/16.20/16.48 | 279.1685 | C_15_H_23_O_3_N_2_ | -1.739 | MS^2^[279]:261;233;205;149;132;120; MS^3^[279→120]:103;93;91 | Isoleucyl-phenylalanine or its isomers |
| 8.73 | 282.1181 | C_11_H_16_O_4_N_5_ | -1.52 | MS^2^[282]:136 MS^3^[282→136]:136;119;94 | 1-Methyladenosine |
| 8.20 | 284.0973 | C_10_H_14_O_5_N_5_ | -1.685 | MS^2^[284]:152 MS^3^[284→152]:152;135;110;109 | Guanosine |
| 26.59 | 290.2670 | C_16_H_36_O_3_N | -1.97 | MS^2^[290]:272;242;122; MS^3^[290→242]:88 | 2-Amino-1,3,4-hexadecanetriol |
| 4.64 | 291.1286 | C_10_H_19_O_6_N_4_ | -1.311 | MS^2^[291]:175  MS^3^[291→175]: 158;157; 130;116;112;70;60 | *N^2^*-(3-Hydroxysuccinoyl)arginine |
| 11.53/12.0/12.37/  13.34 | 295.1635 | C_15_H_23_O_4_N_2_ | -1.724 | MS^2^[295]:277;249;182;165;86;  MS^3^[295→277]: 259;231;166;120 | Isoleucyl-tyrosine or its isomers |
| 11.66 | 297.1541 | C_18_H_21_O_2_N_2_ | -5.614 | MS^2^[297]:265;248  MS^3^[297→265]: 248;221;204;187;176; 161;112 | Alamaridine |
| 7.52 | 314.0901 | C_13_H_16_O_8_N | 3.047 | MS^2^[314]:296;278;136;97  MS^3^[314→136]:136;119;94 | 4-*O*-β-d-Glucopyranoside-  2,4-benzoxazolediol |
| 26.46 | 318.2982 | C_18_H_40_O_3_N | -2.111 | MS^2^[318]:300;256; MS^3^[318→256] 228;212;102;88 | 2-Amino-1,3,4-octadecanetriol |
| 6.87 | 332.1324 | C_14_H_22_O_8_N | -1.643 | MS^2^[332]: 314;233;170;152;136;108 | 5'-*O*-beta-d-Glucosylpyridoxine |
| 1.48 | 337.1701 | C_12_H_25_O_7_N_4_ | -1.706 | MS^2^[337]:319;301;283;260;257;217;209;175;173;158;112 MS^3^[337→319]: 301; 283; 275;260;257; 239;209; 175; 158;112 | *N^2^*-Fructopyranosylarginine |
| 19.43 | 348.1784 | C_19_H_26_O_5_N | -2.139 | MS^2^ [348]: 207;175;142; 122;  MS^3^[348→142]:142;124;122;96;70 | 1,2-Dihydro-*O*-methyltazettine |
| 9.69 | 418.1685 | C_18_H_28_O_10_N | -2.282 | MS^2^[418]: 400;286;238;148 MS^3^[418→238]:220;208;202; 190; 174;172;164; 148;146;134;118;108;106 | Passicapsin |
| 16.78 | 438.2359 | C_25_H_32_O_4_N_3_ | -2.863 | MS^2^[438]:421;292;275;218;204;147; MS^3^[438→204]:147 | Meefarnine B |
| 9.16/9.51 | 454.1685 | C_21_H_28_O_10_N | -2.282 | MS^2^[454]:322;160  MS^3^[454→322]:304;160;142 | *O*-(tri-*O*-Acetyl-α-l-rhamnopyranoside)-(4-hydroxybenzyl) methylcarbamic acid |
| 16.05 | 454.2301 | C_26_H_32_NO_6_ | 7.686 | MS^2^[454]: 437;308;292;275;234;220;204;163;147 | Methyllagerine *N*-oxide |
